# Seasonality dynamics of avian influenza occurrences in Central and West Africa

**DOI:** 10.1101/007740

**Authors:** T. L. Fuller, M. F. Ducatez, K. Y. Njabo, E. Couacy-Hymann, T. Chasar, G. L. Aplogan, S. Lao, F. Awoume, A. Téhou, Q. Langeois, S. Krauss, T. B. Smith

## Abstract

Understanding seasonal cycles of viruses originating in wildlife can provide insight into their likelihood of persistence in animal populations and inform policies to limit spillover to human populations. Avian influenza virus (AIV) is an important zoonotic agent causing seasonal occurrence of avian influenza (AI) in wild birds in temperate areas. Although the seasonality of AIV transmission in tropical birds is largely unknown, peaks of influenza activity in human populations in the tropics coincide with the rainy season. To assess the seasonality of AI in tropical birds, from 2010-14, we sampled 40,099 birds at 32 sites in Central Africa (Cameroon, Central African Republic, Congo-Brazzaville, and Gabon) and West Africa (Benin, Côte d’Ivoire, and Togo). Although AIV was not isolated by egg culture, in Central Africa, detection rates by real-time RT-PCR were 3.57% for passerine songbirds and 8.74% for Anatid ducks. RT-PCR positivity in resident birds increased when Palearctic migrants arrived in Central Africa. At sampling sites with two annual wet seasons, the positive rate in wild birds was greatest during the big rainy season in September – October. This study provides the first evidence that AI is present in Central African birds and identifies environmental factors associated with cases.

## INTRODUCTION

Avian influenza virus (AIV) is an important zoonotic pathogen, resulting in global human morbidity and mortality. Over the past 10 years, poultry and wild birds have transmitted three subtypes of AIV to humans, each posing a substantial health and economic threat (for direct transmission to humans from wild birds, see [1]). Subtype H5N1 has caused 665 human cases in 15 countries in Asia, with a 59% mortality rate [2, 3]. Subtype H7N9, first isolated in 2013, has spread rapidly in East Asia, resulting in 447 human cases with 35% mortality [4, 5]. Subtype H10N8, also isolated in Eastern China in 2013, has caused only two deaths, but could result in a substantial number of human cases if it continues to circulate [6].

A crucial aspect of policies that aim to control AIV transmission to humans is identifying avian reservoirs of the virus. This is especially important in tropical countries where H5N1 has been isolated from birds but the capacity for sampling and screening is typically limited. A region in which the need for surveillance is particularly great is tropical Africa, where H5N1 has been confirmed in eleven countries [7, 8]. To date, surveillance in the region has focused on poultry, ducks, and shore birds [9–12]; however, the prevalence of AIV in other birds merits investigation in light of the discovery that songbirds in the order Passeriformes are also important hosts of H7N9 [5, 13, 14]. For example, sequence analysis of H7N9 viruses isolated from humans indicates that 75% of the genome is genetically similar to that of viruses isolated from songbirds [5]. Furthermore, H7N9 has been isolated from a sparrow in Shanghai, the epicenter of the epidemic [13]. Moreover, transmission experiments demonstrate that songbirds, such as sparrows and finches, infected with H7N9 can spread the virus to other birds via oropharyngeal shedding [14]. Surveillance of African waterfowl has shown that AIV is transmitted from migratory to resident ducks [11], however, the possibility of migratory passerines introducing the virus to resident populations in their wintering areas is unknown. Because surveillance reports of passerines in tropical Africa have been limited to fewer than 300 individuals [9, 12, 15], more extensive sampling is needed to estimate AIV prevalence in African songbirds and assess migratory-to-resident transmission in this avian order.

In addition to determining which avian species are serving as AIV reservoirs in tropical Africa, it is critical to characterize ecological drivers, such as climate patterns, of AIV emergence and zoonotic spillover. In human populations in the tropics, annual influenza epidemics coincide with the wet season, likely because people crowd indoors to avoid rain, increasing the opportunity for virus transmission [reviewed in 16]. However, the question remains as to whether increases in AIV prevalence in tropical birds are also associated with seasonal rains.

The specific objectives of this study were to: 1) determine the prevalence of AI in tropical birds in Central and West Africa, 2) examine the relationship, if any, between rainy season timing and AI occurrences in birds, and 3) assess the evidence for migratory-to-resident bird AIV transmission by sampling resident birds during the season when migrants were present in wintering areas in Central and West Africa.

## METHODS

### Study sites and specimen collection

#### Central Africa

Domestic and wild birds were sampled at 18 sites in Cameroon, Central African Republic, Congo-Brazzaville, and Gabon from October 2010 to July 2013 (Table 1, Fig. 1). Because previous sampling in tropical Africa targeted migratory waterfowl and shorebirds, we primarily screened wild resident songbirds, with opportunistic sampling of chickens and domestic ducks.

**Table 1.**
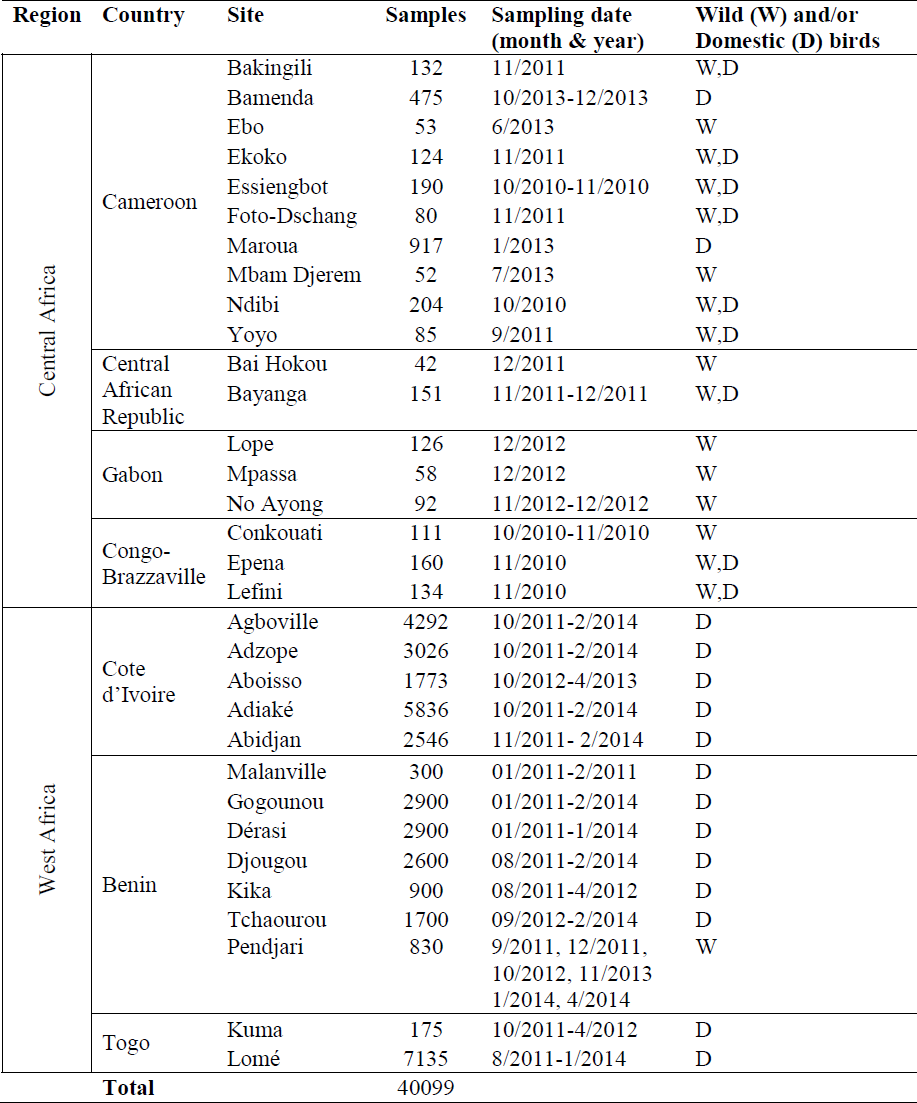
Avian influenza surveillance sites in Central and Western Africa (2010-2014) showing the number of cloacal and oropharyngeal samples per site and the timing of data collection.

**Fig. 1.**
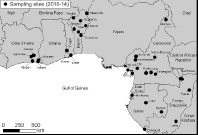
Location of avian influenza surveillance sites in Central and West Africa.

Domestic birds were sampled in village compounds, surrounding farms, and live bird markets. We determined the density of poultry at each site using data reported to the UN Food and Agriculture Organisation (Table S1). Wild birds were captured using mist-nets along agricultural edges and natural habitats near villages. We determined the type of wild bird habitat at each location (e.g., cropland, evergreen forest, or wetland) using a map based on remote sensing (Table S1). Although most sampling occurred in the fall season when Palearctic migrants were present in Central Africa, 85% of the wild bird species sampled were non-migratory. The study region encompasses a wide diversity of rainfall regimes. For example, northern Cameroon is arid, southwestern Cameroon has a single wet season with peak rainfall in July-September, and central Cameroon, Central African Republic, Congo-Brazzaville, and Gabon experience a bimodal rainy season with peaks in May and September-October (Fig. 1, Table S1, Figs. S1-2).

#### West Africa

We sampled 14 sites in Benin, Côte d’Ivoire, and Togo from January 2011 to April 2014. Both cloacal and oropharyngeal swabs were collected from domestic poultry; fecal (environmental), and/or cloacal and oropharyngeal swabs were collected from wild birds in Northern Benin. The domestic birds comprised chickens, guinea fowls, and a few ducks, pigeons and turkeys. The wild birds were primarily passerines.

### Sample processing and molecular analysis

All samples were placed immediately on ice after collection, then stored in liquid nitrogen at -196°C or at -70°C until processed. Côte d’Ivoire samples were screened in the LANADA in Bingerville (Côte d’Ivoire), Benin and Togo samples were either screened on site (in the LADISERO of Parakou, Benin or in the Laboratoire vétérinaire central de Lomé, Lomé, Togo) or in UMR 1225 in Toulouse (France). Central African samples were screened at St. Jude Children’s Research Hospital.

### RNA extraction

Total RNA was extracted from individual swabs using the RNeasy Mini Kit or with the QIAmp Viral RNA Mini Kit (Qiagen, Valencia, CA, USA) following the manufacturer’s guidelines.

### Molecular testing

Swab samples from West Africa were tested as previously described [10]. For the samples from Central Africa, real-time reverse transcription PCR (qRT-PCR) was used to detect the presence of influenza virus genetic material. Viral RNA was amplified using 4x Taqman Fast Virus 1-Step Master Mix, including a universal influenza A forward primer (5’-GACCRATCCTGTCACCTCTGAC-3’), reverse primer (5’-AGGGCATTYTGGACAAAKCGTCTA-3’), and probe (5’-FAM-TGCAGTCCTCGCTCACTGGGCACG-TAMRA-3’) under the following cycling conditions: 1 cycle at 50°C for 5 min; 1 cycle at 95°C for 20 sec; 40 cycles at 95°C for 20 sec and 60°C for 30 sec. Samples showing a cycle threshold (CT) < 40 were considered positive by qRT-PCR and were selected for egg culture. Influenza virus A/California/04/2009(H1N1) was used as a positive control throughout all stages of sample processing.

### Confirmatory rescreening by qRT-PCR

To further assess the evidence for AIV we also carried out a separate round of qRT-PCR screening using a different set of primers. In this round, we rescreened the samples collected in Central Africa in 2010 (*n* = 817) with an additional influenza assay using primers known to amplify a conserved segment of the Matrix I gene of AIV strains that circulate in passerines [17]: 5’-GARATCGCGCAGARACTTGA-3’ and 5’-CACTGGGCACGGTGAGC-3’ were the forward and reverse primers, respectively. In addition, we attempted to amplify and sequence the second subunit of the AIV HA gene using primers HA-1144 [18]: 5’ GGAATGATAGATGGNTGGTAYGG 3′ and Bm-NS-890R [19]: 5’ ATATCGTCTCGTATTAGTAGAAACAAGGGTGTTTT 3′.

### Egg culture

Samples positive by qRT-PCR were subsequently grown in the allantoic cavities of 10-day-old embryonated chicken eggs in attempt to isolate the virus, following established protocols [19].

### Computational analysis

We used a regression model to test for associations between the occurrence of AIV in avian samples and four variables: 1) the avian family to which the sampled species belonged, 2) the percent migratory birds captured at the site, 3) the percent relative humidity at the site, and 4) the number of rainy seasons at the site (Table S1).

#### Avian family

We queried the World Bird Database (AVIBASE) to find the avian family of each bird sampled (Table S1).

#### Migratory birds (%)

We also consulted AVIBASE to determine the migratory status of each bird species; this allowed us to calculate the percent migrants per site.

#### Relativity humidity (%)

In lab settings, the transmission of influenza A is greater in guinea pigs under conditions of low relative humidity [20]. However, surveillance reports in East Africa have detected high AIV prevalence in birds during periods of high relative humidity [21]. We included relative humidity to test whether rates of AIV in Central African birds were associated with high or low relative humidity. The efficiency of influenza A transmission has also been linked to absolute humidity [22], but spatial data for on absolute humidity were not available for the study sites.

#### Number of rainy seasons per site

Rainfall data was obtained from remote sensing as our sampling took place in isolated areas distant from weather stations on the ground. We utilized precipitation radar data from the Tropical Rainfall Measuring Mission (TRMM) satellite, which is routinely used to characterize the volume of rain at ground-level in tropical regions (Table S1). TRMM estimates of average monthly precipitation at each sampling location were validated by comparison with climate maps developed by the United Nations Food and Agriculture Organisation and the Ministry of Agriculture and Rural Development of Cameroon. Based on the precipitation data, we classified each of the sampling locations as having one or two annual wet seasons (Figs. S1-2).

Because we collected multiple samples at each study site, the samples were not statistically independent. In light of this, we required a regression model that would account for the nesting of samples within sites, in order to avoid unacceptably high rates of Type I error in our hypothesis tests. Generalized linear mixed models (GLMM) are typically used in the AIV surveillance literature to account for this issue of pseudo-replication [11]; therefore, we fit a GLMM to our data using PROC GLMMIX in SAS 9.4.

## RESULTS

### Regional comparison

The species composition and abundance of bird communities in Central and West Africa were significantly different (Mantel test *r* = 0.513, p = 1.99 × 10^-4^, for description of the Mantel test, see [23]). For example, 55% of the wild birds sampled in West Africa were not collected in Central Africa. In light of this, we analyzed the two regions separately.

### West Africa

All the Western African samples were negative for AIV by RT-PCR.

### Central Africa

In total, 3.57% of passerine birds and 8.74% of Anatid ducks were positive for AIV by qRT-PCR (using the primers listed in the “Molecular Testing” section of the Methods). There was significant variation in the rate of qRT-PCR positives among avian families (Table 2), driven by the difference in the positive rate between chickens and Anatid ducks. However, there was no statistical difference in the positive rate between Anatids and Passeriformes (Fig. 2).

**Fig. 2.**
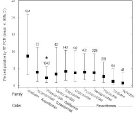
Passerine birds have AIV positive rates similar to Anatid ducks in Central Africa. Only Phasianidae (chickens) had a lower positive rate than Anatids by a *t*-test with a Holm adjustment for multiple comparisons (* indicates *p* = 0.0043). The sample size for each family is listed above the confidence intervals. The plot includes families with 40 or individuals sampled, which comprised 92% of the Central African samples. For families with less than 40 individuals, the number of samples was insufficient to accurately calculate confidence intervals.

None of the attempts to isolate virus in chicken eggs were successful. Because the amount of RNA extracted from cloacal swabs is generally low, without the ability to grow the virus, it was not possible to further characterize AIV subtype via molecular analyses. In order to address the potential of false positive results from an overly sensitive assay, we compared the CT-values of our positive samples to CT-cutoff values typically used in influenza literature and found that our results were significantly lower (one sample t-test *t* = -3.5307, *df* = 339, *p* = 2.36 × 10^-4^; Table S2). While the low CT-values of the positive samples provide relative confidence in the presence of AIV in the original samples, the use of the primers listed in the “Molecular Testing” section of the Methods detected the same rate of AIV positives as the primers listed in the “Confirmatory Rescreening” section (13.3% vs. 10.1%, χ^2^ = 0.1261, *df* = 1, *p* = 0.723). The consistency of the results based on two sets of primers provides additional support that AIV was present in the Central Africa samples. However, elucidating why AIVs circulating in passerines are refractory to growth in egg culture remains an important area for future research.

When we fit the GLMM to the Central African samples, the relationship between relative humidity and qRT-PCR positives approached significance (*p* = 0.0672, Table 2), which is suggestive of a relationship between air moisture and the spread of AIV among tropical birds. The assessed positive rate was significantly higher at sites in Central Africa with a single rainy season than among birds at sites with two rainy seasons (Fig. 3). In addition, the rate of qRT-PCR positives increased with the percentage of migratory birds captured at the sampling location (Fig. 4).

**Table 2.**
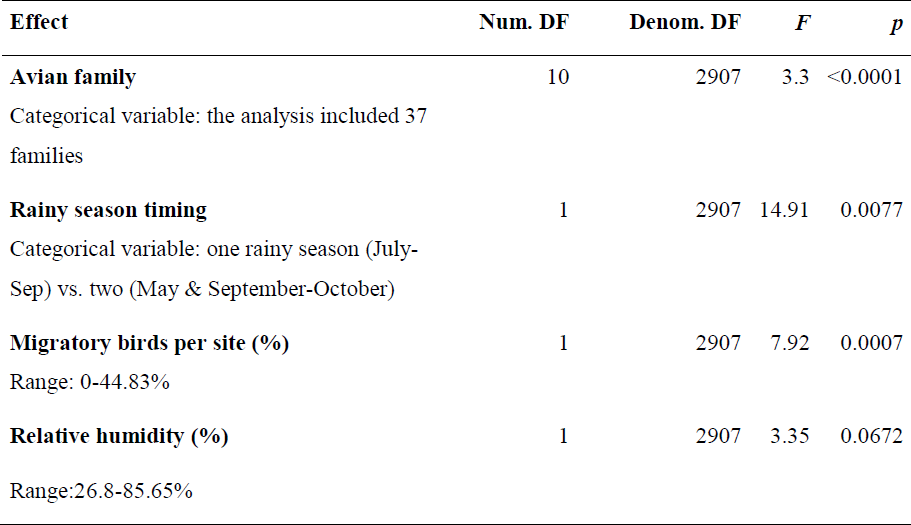
Effect of avian family and ecological variables on AIV positives by RT-PCR in Central African birds. We tested the significance of the predictor variables using a Type 1 Test of Fixed Effects in SAS PROC GLIMMIX.

| Effect | Num. DF | Denom. DF | *F* | *p* |
| --- | --- | --- | --- | --- |
| **Avian family**<br>Categorical variable: the analysis included 37 families | 10 | 2907 | 3.3 | <0.0001 |
| **Rainy season timing**<br>Categorical variable: one rainy season (July-Sep) vs. two (May & September-October) | 1 | 2907 | 14.91 | 0.0077 |
| **Migratory birds per site (%)**<br>Range: 0-44.83% | 1 | 2907 | 7.92 | 0.0007 |
| **Relative humidity (%)**<br>Range:26.8-85.65% | 1 | 2907 | 3.35 | 0.0672 |

**Fig. 3.**
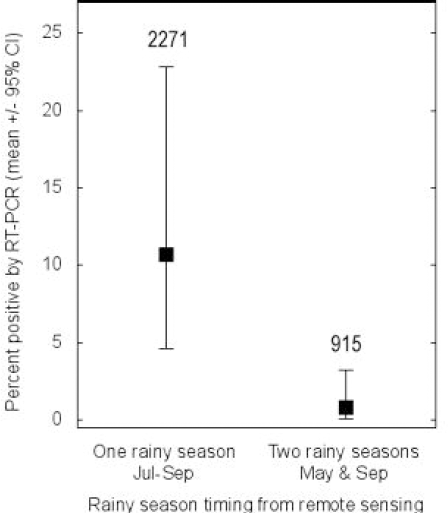
Central African sites where there is a single rainy season have the highest rate of RT-PCR positives (*t*-test with a Holm adjustment for multiple comparisons, *p* = 0.0005). The sample size for each type of rain season is listed above the confidence intervals.

**Fig. 4.**
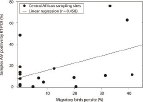
AIV cases in Central African birds increase with the percent migrants at sampling sites. According the linear regression model, when 30% of the birds are migrants, 29% of the samples are AIV positive, whereas when 40% are migrants, 36% are positive.

Since the arrival of Palearctic migratory songbirds and the big rainy season occur simultaneously in September-October in Central Africa, the observation of higher prevalence in areas with a single rainy season may be confounded by the presence of migratory birds. Because the data for percent migrants is continuous and rainy season timing is categorical, there is no linear correlation between the two variables that could be measured with Pearson’s *r* test. Therefore, to assess the relative importance of these variables, we compared the likelihood of models that included rainfall and percent migrants to the likelihood of models that included only one of the variables (Table S3). The two variables had similar weights based on Akaike’s information criterion (Table S4), suggesting that they are equally important drivers of AIV prevalence in Central African birds.

## DISCUSSION

Consistent with previous surveillance reports of West Africa [10], our sampling did not detect AIV in the region’s poultry or wild birds. For the remainder of the Discussion, we focus on Central African countries. Although AIV was not isolated by culture in chicken eggs, the qRT-PCR assay utilized here is highly sensitive and replicable [21, 24]. Indeed, our estimates of AIV positivity by qRT-PCR are consistent with results of previous surveillance efforts in sub-Saharan Africa. A dozen passerine families that we sampled were also screened in a study in southern Africa, and our positive rate estimates for these families were the same as the reported rates in [15] (paired t-test *t* = -1.6, *df* = 11, *p* = 0.14). Furthermore, our 8.74% positive rate for Anatids is comparable with reported rates for African ducks ranging from 5-20% [11, 12, 25]. Our finding that the timing of the rainy season was a significant predictor of the percentage of AIV positives was also supported by surveillance studies in southern Africa [15], which concluded that precipitation events trigger avian dispersal, and as a result spreads AIV among wild bird populations. In addition, surveillance of poultry and wild birds in Uganda found a higher percentage of AIV-positive samples by qRT-PCR during the period of the year that coincided with the wet season and high relative humidity [21]. Similar to the Uganda study, we found that the rate of qRT-PCR positives in Central Africa was greatest during the rainy season, when relative humidity was high (65-95%).

What ecological mechanisms might explain the significant relationship that we observe between AIV positives and timing of rains? Most of our sampling occurred in November, shortly after the beginning of the fall rainy season. The onset of rainfall could explain increases in AIV infections in tropical birds during the wet season via two interrelated mechanisms. The first involves the relationship between rainfall, resource availability, and avian abundance. In tropical rainforests in Australia, rainfall during the wet season leads to increased fruiting, flowering, and abundance of nectar, fruits, and insects; in turn, the increased availability of these food resources results in higher bird abundance [26]. We hypothesize that in Central Africa the rainy season is associated with increased food resources for birds, which leads to higher avian abundance, and increased AIV transmission from infected to susceptible birds. The fall rainy season also coincides with the arrival of Palearctic migratory birds in Central Africa, further increasing avian abundance and providing opportunity for the virus to be transmitted in habitats with high densities of migrants and residents. Several songbird species that were abundant in our sampling sites nest or forage in flocks, which could provide opportunities for AIV transmission within groups, such as the Village Weaver *Ploceus cucullatus* (83 individuals sampled), which forms large colonies with many nests per tree and the Common Bulbul *Pycnonotus barbatus* (53 individuals sampled), which typically forages in groups.

The second mechanism that could explain the occurrence of AIV in tropical birds during the wet season is that seasonal breeders reproduce during this period. Breeding activity in tropical birds is often linked to rainy season onset [27, 28], with the timing of reproduction depending upon the species’ feeding ecology. For example, in Nigeria, insectivores breed at the start of the rainy season when insect abundance is greatest whereas in Nigeria and Cameroon, granivores breed several weeks after rainy season onset, when seeds are at their peak [29]. We hypothesize that AIV is endemic in resident birds in Central Africa, but the prevalence is low for most of the year. When precipitation cues birds to congregate and breed, the resulting increase in contact between infected and susceptible individuals results in a seasonal peak in prevalence.

The rate of AIV positives was higher at locations with a single wet season than at those with a bimodal distribution of rainfall. In comparison with areas with one rainy season, areas with two have a shorter dry season and likely experience less avian mortality due to starvation during food-lean times. As a result, peaks in bird abundance are likely lower in areas with two rainy seasons and hence there is less inter-annual variation in AIV prevalence. Furthermore, the environmental cue to congregate and breed could be weaker in regions where the breeding season may be more protracted.

Sites with one or two rainy season(s) may also differ with respect to other factors that could impact the positive rate, such as habitat type, avian density, and the species composition of bird communities. This raises the possibility that the effect of the number of rainy seasons on AIV prevalence, while statistically significant, may be an artifact of other factors that differed between our sampling locations. To assess this, we tested for differences in poultry density between sites with one and two-rainy seasons using a *t*-test, tested for differences in habitat types between single and dual rainy season sites using a chi-squared test, and tested for differences in species composition using a Mantel test. When we compared sites with a single rainy season to those with two rainy seasons, there was no difference in poultry density (*t* = 0.8553, df = 5.981, *p* = 0.415), habitat types (χ^2^ = 4.5, df = 5, *p* = 0.4799), or wild bird species composition (Mantel test *r* = 0.0668, *p* = 0.192). This suggests that the significant association between qRT-PCR positives and the number of rainy seasons was not merely an artifact of other habitat factors.

We found evidence for the transmission of AIV from migratory to resident songbirds to the extent that the rate of positives increased with the percentage of migrants captured at the sampling location. This is consistent with water bird surveillance in Africa, which has shown that AIV prevalence increases with the arrival Palearctic migrants in September [11, 25, 30]. Our statistical model suggests that the timing of the rainy season is an important driver of AI in songbirds independent of the fact that the fall rainy season coincides with Palearctic migration (Tables S3-4). However, disentangling the relative importance of rainy season timing and the migrant-to-resident spillover is clearly needed.

We hypothesize that the wet season triggers increased bird abundance and breeding, increasing contact between infectious and susceptible individuals and resulting in a seasonal peak in prevalence. An alternative hypothesis that could be tested by sampling during the dry season is that wild birds congregate around rare water sources, a pattern observed in Nigeria, where AIV prevalence peaks in the dry season because the disappearance of small ponds results in increased waterfowl density at remaining ponds, leading to increased AIV transmission [31]. If this also occurs in Central Africa, sampling during the dry season might detect a high percentage of AIV positives.

To control for the effect of Palearctic migrants, one could sample resident birds during the rainy season but prior to the arrival of migrants in Central Africa. If spillover from migrants were a more important driver of AIV than the timing of rainfall, one would expect lower prevalence at sites sampled before the migratory period. As only 5.81% of our Central African samples were collected in the spring or summer prior to the arrival of migrants (Figs. S2-S3), additional sampling is required to make robust inferences about spring versus fall prevalence. The collection of additional spring samples will also clarify the relative importance of the wet season and the arrival of Palearctic migrants as drivers of AI in Central African birds.

The present analysis contributes to the understanding of AIV circulation in wild birds by confirming surveillance reports which have found no AIV in West Africa [10] and sampling Central African Republic, Congo-Brazzaville, and Gabon in Central Africa, which are countries that have not previously been surveyed. The positive rate that we detected in Central Africa’s wild birds is consistent with surveillance elsewhere in the Old World Tropics. For example, infection rates in passerines at tropical sites in northern Vietnam and southern China range from 2.3-6.6% [32, 33], which is within the confidence limits of our estimates for Central Africa (1.57-7.95%).

This analysis also broadens our knowledge of AIV dynamics in wild birds by providing the first evidence that infections are seasonal in songbirds in equatorial Africa. It has long been appreciated that in temperate areas, wild bird populations experience seasonal peaks in AIV prevalence. These include autumn peaks in mallards in Alberta, Canada [34] and Öland, Sweden [35] and spring peaks in shorebirds at Delaware Bay, US [36]. Factors hypothesized to trigger this seasonality in temperate birds include prey abundance, avian behavioral and physiological cycles, the pulse of births in the spring, and migratory and hormonal cycles [36, 37].

The detection of AIV by RT-PCR in the present study suggests the possibility of that the virus circulates widely in Central Africa birds, a finding that should stimulate further surveillance to isolate the virus by egg culture. Expanding AIV screening programs in Central and West Africa can confirm the qRT-PCR positives reported here and possibly obtain isolates for molecular characterization and pathogenicity studies. If the AIV subtypes circulating in birds in tropical Africa were low pathogenic with a putative low pandemic risk (as could be assessed using risk assessment tools as described in [38]), controlling the spread of the virus may not be urgent. However, if highly pathogenic subtypes H5 or H7 occur in the region, then shifts in poultry rearing practices would be warranted to limit spillover from wild birds to domestic animals and humans. Since subtype H5N1 has already caused losses of $20 billion to the global poultry industry [39], it could have a substantial impact on food production in Central and West Africa. Furthermore, subtype H7N9, which has high virulence in humans and circulates in songbirds in China, could result in treatment costs of $5.3 billion if it spreads to a major city [40]. Improving our understanding of the occurrence and seasonality of AIV in African birds can provide insights useful for the formulation agricultural and biosecurity policies.

## SUPPLEMENTARY MATERIAL

For supplementary material accompanying this paper visit URL TO BE ASSIGNED UPON ACCEPTANCE

## ACKNOWLEDGEMENTS

We thank Nicole Arrigo, Richard Webby, and Simon Ducatez for comments on the manuscript. We also acknowledge Viviane Kouakou, Yapi Yapo, Danho Thérèse, Koffi Yao Mathurin, Gnabro Privat, Kouassi Sue Lou Antoinette, Nana Pauline, Agolai Innocent,Toussaint Lougbégnon, Tawaliou Alidou, Assana Garba Bankole, Tatiana Toure, Séverin Adjitore, Jeanne Abdoulaye, Rodrigue Setchegbe, Adimi Adje Sylvain, Alao Funmi, Lengo Kossiwa, Go-Maro Wolali, Dogno Koffi, Pali Magnoudéwa, Kpatina Alfred, Voedjo Koukpealedou, Aketre Yawo, Charlotte Foret, Angélique Teillaud, Josyane Loupias, Brigitte Peralta, Charley Lagarde, Christelle Camus-Bouclainville, Guillaume Croville, Etienne Liais, Renaud Berger, Clément Fage, Florian Grard, Jean-Benoit Tanis, Brittany Hale, and Manon Tournou for their excellent technical assistance. We also thank Amanda Ball, James Knowles, Jennifer DeBeauchamp, Jerry Parker, Richard Elia, Pamela McKenzie, and Maureen Rice for help with logistics and data management.

